# Assessment of developmental neurotoxicology-associated alterations in neuronal architecture and function using *Caenorhabditis elegans*

**DOI:** 10.1101/2025.01.11.632560

**Authors:** Javier Huayta, Sarah Seay, Joseph Laster, Nelson A. Rivera, Abigail S. Joyce, P. Lee Ferguson, Heileen Hsu-Kim, Joel N. Meyer

**Affiliations:** Nicholas School of the Environment, Duke University, Durham, North Carolina, USA; Pratt School of Engineering, Duke University, Durham, North Carolina, USA

**Keywords:** Developmental neurotoxicity, DNT, neurodegeneration, altered behavior, *Caenorhabditis elegans*

## Abstract

Few of the many chemicals that regulatory agencies are charged with assessing for risk have been carefully tested for developmental neurotoxicity (DNT). To speed up testing efforts, as well as to reduce the use of vertebrate animals, great effort is being devoted to alternate laboratory models for testing DNT. A major mechanism of DNT is altered neuronal architecture resulting from chemical exposure during neurodevelopment. *Caenorhabditis elegans* is a nematode that has been extensively studied by neurobiologists and developmental biologists, and to a lesser extent by neurotoxicologists. The developmental trajectory of the nervous system in *C. elegans* is easily visualized, normally entirely invariant, and fully mapped. Therefore, we hypothesized that *C. elegans* could be a powerful *in vivo* model to test chemicals for the potential to alter developmental patterning of neuronal architecture. To test whether this might be true, we developed a novel *C. elegans* DNT testing paradigm that includes exposure throughout development, examines all major neurotransmitter neuronal types for architectural alterations, and tests behaviors specific to dopaminergic, cholinergic, and glutamatergic functions. We used this paradigm to characterize the effects of early-life exposures to the developmental neurotoxicants lead, cadmium, and benzo(a)pyrene (BaP) on dopaminergic, cholinergic, and glutamatergic architecture. We also assessed whether exposures would alter neuronal specification as assessed by expression of reporter genes diagnostic of specific neurotransmitters. We identified no cases in which the apparent neurotransmitter type of the neurons we examined changed, but many in which neuronal morphology was altered. We also found that neuron-specific behaviors were altered during *C. elegans* mid-adulthood for populations with measured morphological neurodegeneration in earlier stages. The functional changes were consistent with the morphological changes we observed in terms of type of neuron affected. We identified changes consistent with those reported in the mammalian DNT literature, strengthening the case for *C. elegans* as a DNT model, and made novel observations that should be followed up in future studies.

## 1. Introduction

Environmental exposure to toxicants during development can result in DNT, but very few chemicals have been tested carefully for DNT (Crofton et al., 2012; Mundy et al., 2015; Raffaele et al., 2010). Therefore, great effort is being devoted to development of new approaches for testing DNT (Coecke et al., 2007; Fritsche et al., 2018; Lein et al., 2007; Li et al., 2017; Sachana et al., 2019), with increasing use of non-vertebrate models required by law in both the United States and the European Union (Bal-Price and Fritsche, 2018; Moné et al., 2020). DNT is an outcome from several mechanisms, including alterations in cell fate, cell structure, and neuronal function (Tamm and Ceccatelli, 2017). Here, we focus on changes in cell structure (“architecture”), and neuronal function. Vertebrate models are closer physiologically to humans than invertebrate models are. However, assessment of alterations in cell architecture can be challenging in vertebrates. Most vertebrate models are not transparent, so assessment of neuronal architecture is more complicated, typically involving histological experiments. Even in those that are transparent (e.g., fish strains with pigment genes mutated), neuronal populations and structures are not invariant, complicating assessment of alterations to those patterns (Miller et al., 2018). On the other hand, *in vitro*, cellular models are being developed, and some permit interrogation of changes in growth of cell projections and other *in vitro* measures of changes to cell architecture (Barbosa et al., 2015; Breier et al., 2010). We hypothesized that *C. elegans* could provide a useful, intermediate complement to vertebrate and cell culture models. Reasons include: 1) *C. elegans* is transparent, such that visualization of neuronal identity and structure is easy, especially with transgenic reporter strains; 2) normally, neuronal fate and morphology are invariant (302 neurons and 56 glial cells), and synaptogenesis is nearly invariant, such that any deviation is easily observed; 3) the developmental trajectory and neurotransmitter identities of the *C. elegans* neurons are exceptionally well-described due to decades of work by *C. elegans* developmental biologists and neurobiologists; 4) the processes of neuronal cell fate specification are in many cases conserved with those observed in mammals and can be perturbed genetically (Hobert, 2010), suggesting that environmental perturbation of those processes should be testable in *C. elegans*, with potential extrapolation to vertebrates; 5) *C. elegans* offers a wide range of other advantages for toxicological studies, including the ability to carry out experiments cost-effectively with a large number of biological and technical replicates and large number of exposure concentrations, assessment of different life stages due to a short lifespan with well-conserved biological hallmarks of aging, and other advantages, as we (Hartman et al., 2021; Leung et al., 2008; Maurer et al., 2018, 2015; Weinhouse et al., 2018) and others (Hunt, 2017; Meyer and Williams, 2014; Peterson et al., 2008; Ruszkiewicz et al., 2018; Tejeda-Benitez and Olivero-Verbel, 2016) have reviewed. Despite these apparent advantages, *C. elegans* has been used in only a limited fashion for DNT *per se*, with a focus to date on behavioral outcomes (Boyd et al., 2010; Hunt et al., 2018; Ijomone et al., 2020).

However, the *C. elegans* nervous system is simpler than that of larger, more complex animals. For example, *C. elegans* transmits information using non-spiking neurons, and has only four pairs of olfactory amphid neurons with a broad range of odorant-specificity compared to neurons with odorant-specificity in mammals, indicating a high level of sensory integration (Naudin et al., 2022; Schafer, 2016). In addition, while many of the mechanisms regulating neurodevelopment in *C. elegans* are conserved in people, not all are (Hobert, 2010). Therefore, *C. elegans* may offer important advantages in DNT testing, but it is also possible that differences in the development, structure, and function of the worm’s nervous system limits extrapolation of DNT results to people. To begin to assess whether *C. elegans* is likely to be a good model for testing chemical DNT mediated by alterations in neuronal architecture, we tested whether the well-studied human DNT chemicals lead (Pb), cadmium (Cd), and BaP would alter multiple neuronal types, based on previous literature for Pb (Basha et al., 2012; Devi et al., 2005; Lu et al., 2018; Reddy et al., 2007; Tang et al., 2019; Zhou et al., 2000), Cd (Antonio et al., 2010; Chow et al., 2008; Lu et al., 2018; Rai et al., 2010; Tang et al., 2019), and BaP (Gao et al., 2017; He et al., 2012; McCallister et al., 2008; Slotkin et al., 2017, 2013). We looked for changes to neuronal fate (type), altered morphology, and altered function. Because DNT can manifest early or later in life (EPA, 1998), and may present only upon subsequent stressor challenge, we assessed alterations both during development and later in life with a secondary challenge and behavioral assessment.

We found that Pb, Cd, and BaP caused DNT in dopaminergic, cholinergic, and glutamatergic neurons reflected in alteration of dendrite morphology of neurons located in the head of *C. elegans*, and in late-life behaviors associated with these neuronal types, without detectable changes to neuronal cell fate.

## 2. Materials and methods

### 2.1. *C. elegans* strains and maintenance

*C. elegans* strains BY200 (*vtIs1*[p*dat-1*::GFP]), DA1240 (*adIs1240*[*eat-4*::sGFP+*lin-15*(+)]), LX929 (*vsIs48*[*unc-17*::GFP]), CZ1632 (*juIs76*[*unc-25*p::GFP+*lin-15*(+)]), GR1366 (*mgIs42*[*tph-1*::GFP+*rol-6*(su1006)]), VT1485 (*maIs188*[*mir-228*p::GFP+*unc-119*(+)], CB1112 (*cat-2*(e1112) II), CB1141 (*cat-4*(e1141) V), NM1657 (*unc-10*(md1117) X), VC223 (*tom-1*(ok285) I), CB75 (*mec-2*(e75) X), and CB1339 (*mec-4*(e1339) X) were maintained at 20 °C on K-agar plates (Williams and Dusenbery, 1988) seeded with OP50 *E. coli*. The GFP reporter strains were developed by other laboratories to allow identification of neuronal types based on expression of GFP driven by a promoter specific to each neurotransmitter (**Table 1**).

**Table 1:** *C. elegans* strains used for DNT evaluation in different neuronal types.

| Strain | GFP-tagged gene | Neurotransmitter | Reference |
| --- | --- | --- | --- |
| BY200 | *dat-1* | Dopamine | (Nass et al., 2002) |
| DA1240 | *eat-4* | Glutamate | (Lee et al., 1999) |
| LX929 | *unc-17* | Acetylcholine | (Chase et al., 2004) |
| CZ1632 | *unc-25* | GABA | (Ackley et al., 2005) |
| GR1366 | *tph-1* | Serotonin | (Sze et al., 2000) |
| VT1485 | *mir-228* | Multiple (glial cell) | (Martinez et al., 2008) |

### 2.2. Chemicals

Lead acetate (Sigma-Aldrich) and cadmium chloride (Sigma-Aldrich) stocks were dissolved in water to prepare 20 mM stock solutions. BaP (Sigma-Aldrich) was dissolved in DMSO to prepare a 10 mM stock solution. Aldicarb (Sigma Aldrich) was dissolved in ethanol to prepare a 100mM stock solution. 6-hydroxydopamine (Sigma-Aldrich) was dissolved in water to prepare a 100 mM fresh stock solution before experiments. Chlorpyrifos (Sigma-Adrich) was dissolved in DMSO to prepare a 1 mM stock solution. Quinolinic acid (Sigma-Aldrich) was dissolved in DMSO to prepare a 10 M stock solution. AF64A (ethylcholine mustard aziridinium ion) was generated as previously described (Clement and Colhoun, 1975a, 1975b; Fisher et al., 1982). Briefly, acetylethylcholine mustard HCl (Sigma-Aldrich) was dissolved in water to prepare a 10 mM solution, NaOH 10 N was gradually added until reaching pH 11.5 and left for 20 minutes at room temperature. Concentrated HCl was added until reaching pH 7.5 and the solution was left to stand in room temperature for 1 hour. Stock solutions were preserved at −80 °C, and AF64A was prepared fresh before exposure assays. For the exposure experiments, K+ media was prepared with the following composition: 32 mM KCl, 51 mM NaCl, 13 μM cholesterol, 3 mM CaCl_2_, and 3 mM MgSO_4_, corresponding to approximately 90 mM chloride and expected pH of 6.5 (Boyd et al., 2009; Meyer et al., 2010).

### 2.3. Determination of exposure concentrations

A mixed adult worm population was treated with sodium hypochlorite to recover the progeny and generate an age-synchronized population. These embryos were transferred to 24-well plates containing each 500 µL of K+ medium (Boyd et al., 2009) with HB101 *E. coli* at OD 2.0 (measured at 600 nm), approximately 100 embryos and either 0, 50, 100, 150, 200, or 250 µM lead acetate; 0, 0.5, 5, 10, 25, or 50 µM cadmium chloride; or 0, 0.05, 0.10, 0.30, 0.50, or 1.00 µM BaP.

In K+ media, aqueous Cd and Pb were primarily mixtures of free ion forms (Cd^2+^, Pb^2+^) and complexes with chloride ions, as indicated by equilibrium speciation calculations performed with Visual MINTEQ v.3.1 (Gustafsson, 2020). At Pb levels exceeding 100 µM, the calculations indicated that certain Pb-hydroxy-sulfate minerals were close to or just above saturation (i.e., saturation indices between 0 and +1). While we did not observe precipitates forming, the formation of colloidal Pb was a possibility. Cd minerals were undersaturated at all exposure conditions.

After reaching the L4 stage, worms were collected and transferred to unseeded K-agar plates. Plates containing worms were imaged using a Keyence BZ-X710 microscope using a Nikon 4X objective. Images were analyzed using the WormSizer add-in in ImageJ to determine individual worm length of a minimum of 50 individuals (Moore et al., 2013). Regression analysis to determine EC_10_ values for length was performed using MS Excel 365.

### 2.4. Chemical exposures

Worms were age-synchronized by treating a mixed adult population with sodium hypochlorite to dissolve the bodies of gravid adults and obtain their fertilized embryos as previously described (Stiernagle, 2006). These embryos were transferred to K+ medium and allowed to hatch overnight. L1 larval stage worms were collected the next day and transferred to K-agar plates seeded with OP50 *E. coli* and left for 48 hours to reach their L4 larval stage. These worms were transferred to 24-well plates containing each 500 µL of K+ medium with HB101 *E. coli* at OD 2.0, approximately 100 L4 stage worms, and either 140.0 µM lead acetate, 11.7 µM cadmium chloride, or 0.30 µM BaP. Control wells and those containing BaP also contained 1% DMSO as vehicle. After 24 hours, this parent population was collected by transferring the well content to a 15 mL conical tube. The volume was raised to 15 mL with K-medium. After allowing worms to settle for 5 minutes, the supernatant was vacuumed, and this washing step was repeated two more times. Worms were then suspended in 5 mL of K-medium containing final concentrations of 0.4 N sodium hydroxide and 20% v/v sodium hypochlorite to recover their embryos. The progeny population was transferred to plates with OP50 *E. coli* for 52-54 hours to allow them to reach the L4 stage, when quantification of neurodegeneration was performed.

### 2.5. Quantification of morphological alterations to neurons

When exposed worms reached the L4 stage, worms were washed three times and transferred to a 2% agarose pad on a glass slide. Z-stacks were acquired for individual worms’ heads on a Keyence BZ-X710 microscope equipped with a Keyence BZ-X700E light-source and using a Nixon 40X (air) objective with 200 milliseconds of exposure with the ‘high-sensitivity’ option selected. Maximum projections of the Z-stacks were generated, and each dendrite of the cephalic dopaminergic neurons was scored on a scale of 0 to 6 as previously described (Bijwadia et al., 2021). Briefly, 0 – no damage, 1 – irregular (curves), 2 – less than 5 blebs, 3 – 5 to 10 blebs, 4 – more than 10 blebs and/or breaks, 5 – breaks, 25 to 75% dendrite loss, and 6 – breaks, more than 75% dendrite loss. We developed a similar scoring system for cholinergic neurons: 0 – no damage, 1 – irregular (curves), 2 – 1 to 10 blebs, and 3 – more than 10 blebs and/or presence of breaks; for glutamatergic neurons: 0 – no damage, 1 – 1 to 5 blebs, 2 – 6 to 10 blebs, and 3 – more than 10 blebs and/or presence of breaks; and for serotonergic neurons, GABAergic neurons and glial cells: 0 – no damage, and 1 – any type of damage. Statistical analysis was performed using the chi-squared test with Bonferroni correction of *p* values.

### 2.6. Secondary challenge to neurotoxicant

A mixed adult worm population was treated with sodium hypochlorite to obtain an age-synchronized population as described in the previous section. Worms were grown on K-agar plates with OP50 *E. coli* until reaching day 4 of adulthood. These worms were collected and transferred to 24-well plates containing 500 µL of K+ medium and either 0, 10, 25, 50, 75, or 100 mM 6-hydroxydopamine; 0, 1, 5, 10, 25, or 50 µM chlorpyrifos; 0, 1, 5, 10, 50, or 100 µM AF64A; or 0, 10, 50, 100, or 250 mM quinolinic acid. After one hour, worms were collected and washed three times, then transferred to new plates for 24 hours. Neurodegeneration was quantified as described in the previous section and EC_50_ values were determined for all subsequent secondary challenge experiments. Regression analysis to determine EC_50_ values for neurodegeneration was performed using MS Excel 365.

The progeny of worms exposed to either lead acetate, cadmium chloride, or BaP were allowed to grow until day 4 of adulthood on K-agar plates with OP50 *E. coli*. These worms were collected and transferred to 24-well plates containing 500 µL of K+ medium and either 60.5 mM 6-hydroxydopamine, 13.7 µM chlorpyrifos, 77.0 µM AF64A, or 94.1 mM quinolinic acid. After one hour, worms were collected, washed three times, and transferred to new plates for 24 hours. Neurodegeneration was quantified as described in the previous section.

### 2.7. Assessment of basal slowing response (BSR) for dopaminergic function

The progeny of worms exposed to either lead acetate, cadmium chloride, or BaP were allowed to grow until day 4 of adulthood on K-agar plates with OP50 *E. coli*. At this stage, worms were tested for their basal slowing response as previously described (Sawin et al., 2000). Briefly, assay plates were prepared by spreading HB101 *E. coli* in the shape of a ring on K-agar plates and incubated overnight. Worms were collected, quickly washed, and 20-30 worms were transferred to the center of either an assay plate or a plate with no bacteria; with excess liquid absorbed using a Kimwipe. After 5 minutes, 5 videos of 20 seconds each per plate were acquired using a Keyence BZ-X710 microscope. These videos were used to count the number of body bends in 20 seconds of 10-20 worms per plate.

For the positive control strains CB1112 and CB1141, mixed adult populations were treated with sodium hypochlorite to obtain an age-synchronized population as described in the previous section. These worms were grown until day 4 of adulthood and then tested for basal slowing response.

### 2.8. Assessment of sensitivity to soft touch for glutamatergic function

The progeny of worms exposed to either lead acetate, cadmium chloride, or BaP were allowed to grow until day 4 of adulthood on K-agar plates with OP50 *E. coli*. At this stage, worms were tested for their sensitivity to soft touch as previously described (Chalfie and Sulston, 1981; Griffin et al., 2019). Briefly, worms were tested by stroking their heads and tails consecutively with an eyelash mounted on a Pasteur pipette under a Leica MZ75 stereo microscope. Sensitive worms started backwards movement after touching their heads and stopped or started forward locomotion after touching their tails. This process was repeated 5 times per worm for a total of 20-30 worms per replicate, counting the number of times that a worm showed response to soft touch.

For the positive control strains CB75 and CB1339, mixed adult populations were treated with sodium hypochlorite to obtain an age-synchronized population as described in the previous section. These worms grew until day 4 of adulthood and then tested for their sensitivity to soft touch.

### 2.9. Assessment of aldicarb-induced paralysis for cholinergic function

The progeny of worms exposed to either lead acetate, cadmium chloride, or BaP were allowed to grow until day 4 of adulthood on K-agar plates with OP50 *E. coli*. At this stage, worms were tested for their sensitivity to aldicarb-induced paralysis as previously described (Mahoney et al., 2006). Briefly, assay plates containing 1 mM aldicarb were prepared one day before the assay and stored at 4 °C. Plates were allowed to reach room temperature before starting the assay. A drop of 2-3 µL of OP50 *E. coli* was added to the center of the plate and allowed to dry for 20 minutes. 25-30 worms were added to the bacteria spot on the plate and paralysis was assessed every 30 minutes for 6 hours. For our purposes, a paralyzed worm was defined as not showing movement after prodding with an aluminum wire.

For the positive control strains NM1657 and VC223, mixed adult populations were treated with sodium hypochlorite to obtain an age-synchronized population as described in the previous section. These worms were grown until day 4 of adulthood and then tested for their sensitivity to aldicarb-induced paralysis.

### 2.10. Determination of chemical uptake

We used inductively coupled plasma mass spectrometry (ICP-MS) to quantify the uptake of Pb and Cd into worms. A mixed adult worm population was treated with sodium hypochlorite to obtain an age-synchronized population as described in the previous section. Worms were grown on K-agar plates with OP50 *E. coli* until reaching the L4 larval stage. Worms were collected, washed, and approximately 100 worms were transferred to each well of 24-well plates containing K+ medium, HB101 *E. coli*, and either 0, 70, 140, or 280 µM of lead acetate; or 0, 5.9, 11.7, or 23.4 µM of cadmium chloride. After 24 hours, worms and medium were transferred to 5 mL centrifuge tubes and the worms were pelleted at 600×*g*. The supernatant was removed and saved for analysis. The worms were washed a total of 3 times as quickly as possible by adding 5 mL K-medium and centrifuging at 600×*g*. Following the last spin, the liquid was removed, and the worm pellet was stored at −80 °C. 25 µL of worm or medium sample were mixed with 100µL of nitric acid (trace metal grade, Fisher Scientific). The mixture was heated on a hot block at 95 °C for two hours and then diluted to 0.5 mL after cooling. Aliquots of the digestates were analyzed for Pb and Cd in an Agilent 7900 ICP-MS.

We used gas chromatography-mass spectrometry (GC-MS) to quantify the uptake of BaP into worms. A mixed adult worm population was treated with sodium hypochlorite to obtain an age-synchronized population as described in the previous section. Worms were grown on K-agar plates with OP50 *E. coli* until reaching the L4 larval stage. Worms were collected, washed, and approximately 100 worms were transferred to each well of 24-well plates containing K+ medium, HB101 *E. coli*, and either 0, 0.3, 3, or 10 µM benzo(a)pyrene. After 24 hours, worms and medium were transferred to 5 mL centrifuge tubes and the worms were pelleted at 600×*g*. The supernatant was removed and saved for analysis. The worms were washed a total of 3 times as quickly as possible by adding 5 mL K-medium and centrifuging at 600×*g*. Following the last spin, the liquid was removed, and the worm pellet was stored at −80 °C. Worm pellets and supernatant samples were extracted in hexane (400 mL) via sonication for 15 min in a Brason 5510 ultrasonic cleaner, two times, and combined. Extracts were analyzed by GCMS using an Agilent 7890A gas chromatography system, 5975 mass spectrometer, and 7693 auto samplers.

### 2.11. Determination of embryo viability

A mixed adult worm population was treated with sodium hypochlorite to obtain an age-synchronized population as described in the previous section. Worms were grown on K-agar plates with OP50 *E. coli* until reaching the L4 larval stage. Worms were collected, washed, and approximately 100 worms were transferred to 24-well plates containing 0.5 mL of K+ medium each, HB101 *E. coli* as food source, and either 0, 50, 200, or 500 µM lead acetate; 0, 25, 50, 100, 200, or 500 µM cadmium chloride; or 0, 10, 25, or 50 µM benzo(a)pyrene. After 24 hours, worms were washed and transferred to new plates containing 1 worm per plate for a minimum of 10 worms per treatment. Worms were transferred to new plates every 24 hours for 6 days, and the larvae present in each plate counted.

### 2.12. Statistical analysis

MATLAB R2023a (MATLAB 9.14) and OASIS 2 (Han et al., 2016) were used for all statistical testing and graph generation. Statistical tests are described in their corresponding figure legends.

## 3. Results

### 3.1. Development of a DNT paradigm to test the effects of Pb, Cd, and BaP in *C. elegans*

Most previous developmental exposures in *C. elegans* have been carried out beginning at the first larval stage. However, although some neurodevelopment does occur subsequently, most neurons and synapses are fully formed by this stage (Hobert, 2010; Sulston et al., 1983; Sulston and Horvitz, 1977; Witvliet et al., 2021). Therefore, so that all neurodevelopmental processes will be included, we developed a maternal exposure DNT protocol that ensures exposure of the offspring (F1 generation) throughout embryogenesis. In addition, we wanted to be sure that the exposure concentrations were not too high (in which case might identify nonspecific DNT secondary to generalized maternal or embryonic toxicity) and not too low (which would lead to many false negative results).

To this end, we exposed worms to increasing concentrations of Pb, Cd, and BaP during their development to determine the concentrations of these chemicals leading to a 10% reduction in length (EC_10_) after reaching their last larval stage (**Fig. 1A-C**). We calculated EC_10_ values of 140 µM lead acetate, 11.7 µM cadmium chloride, and 0.30 µM BaP. We also wanted to ensure that the concentrations we used did not have large effects on reproduction, which might confound our assessment of effects in offspring. We had previously tested the effects of BaP on reproduction, and did not see statistically significant effects until 10 µM (Harris et al., 2020). We had also previously tested cadmium exposures (Leuthner et al., 2022), and others have tested cadmium and lead for effects on reproduction (Anderson et al., 2001; Guo et al., 2009). Based on those reports, our growth EC_10_ values for lead, cadmium and benzo(a)pyrene should cause minor or reproductive impairment. However, the exposure paradigms in those reports were not identical to this study. To ensure that results were similar in our hands, we tested cadmium, lead, and benzo(a)pyrene. We found that the total number of viable embryos from worms exposed to cadmium chloride did not change at any tested concentration, significantly decreased for worm exposed to 200 µM or more of lead acetate and decreased for worms exposed to 50 µM BaP (**Fig. S1A, C, E**). However, the highest concentration tested of 500 µM cadmium chloride caused a statistically significant delay in reproduction (shift to the right when graphing the number of viable embryos per day). 50 µM BaP did not cause a statistically significant change when analyzed as viable embryos per day, and lead acetate caused a significant decrease in viable embryos per day at 200 µM or higher concentrations (**Fig. S1B, D, F**). However, the EC_10_ values used for the DNT studies were under the concentrations that would affect reproduction. Therefore, we used the calculated EC_10_ concentrations for all further developmental exposures.

**Figure 1:**
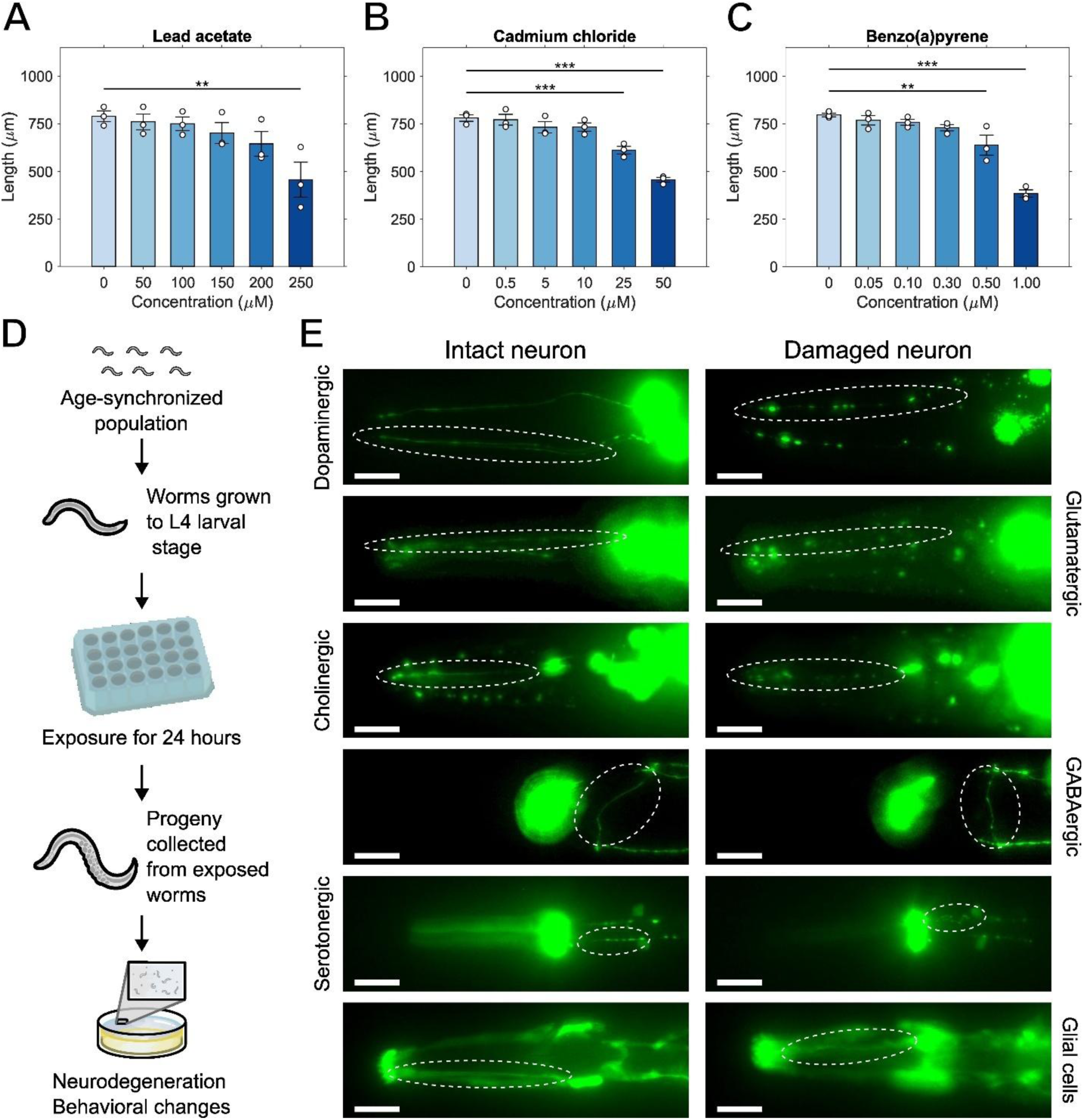
Experimental concentrations and design. Dose response of length measured at the L4 stage of worms developmentally exposed to (A) lead acetate, (B) cadmium chloride, and (C) BaP. (D) Scheme of experimental design. (E) Example images of intact (left) and damaged (right) neurites for all neuronal types. White dashed circles delineate the regions scored. Scale bars 20 µm, One-way ANOVA with Dunnet’s test, (**) *p* < 0.01, (***) *p* < 0.001.

Having established appropriate exposure concentrations, we exposed an age-synchronized parent population when they reached the L4 larval stage when egg production is beginning. After 24 hours of chemical exposure, we recovered the progeny from this parent population for quantification of morphological alterations and changes in behavior (**Fig. 1D**). We identified strains expressing GFP in all neuronal types (as defined by neurotransmitter: **Table 1**), and neurites within those neuronal types that were amenable to consistent scoring. We tested all these neuronal types for the morphological effects of Pb, Cd, and BaP (**Fig. 1E**). We observed various types of irregularities such as kinks, blebs, and loss of dendrite integrity in the dopaminergic, glutamatergic, cholinergic, GABAergic, and serotonergic neurons located in the head, as well as in glial cells located in the same area.

### 3.2. Chemical uptake of Pb, Cd, and BaP in *C. elegans*

The concentrations we used were high relative to cell culture, which is often needed in worm studies because of poor uptake (Hartman et al., 2021). To facilitate comparisons to studies in other models and in people, we next quantified the internal chemical concentrations of Pb, Cd, and BaP after a 24-hour exposure beginning at the L4 larval stage (**Fig. 2**). We found that for lead, concentration in worms was in the range of 11% to 16% of that in the media (**Fig. 2A**) and statistically significantly lower in worms than media for all concentrations tested except the 0 µM control. However, the concentration quantified in the medium corresponds well with that of the original working concentration calculated at the start of the exposure. Interestingly, the internal concentration in worms did not increase proportionally with that in the medium. It is possible that this may result from the formation of colloidal or other non-bioavailable forms of Pb at higher concentrations. In contrast, cadmium internal uptake in exposed worms was not significantly different than the concentration measured in the liquid medium, and in all cases the internal concentration was at least 73% that of the medium (**Fig. 2B**). Furthermore, the concentrations of cadmium in both worms and the liquid medium were a fraction (roughly half) of the original working concentration. Finally, we quantified BaP in both worms and their liquid medium, but concentrations were under the detection limit for all exposed worms (**Fig. 2C**). Notably, for the 3 and 10 µM concentrations, the measured concentrations were ~12% of the original. It is possible that the experimental conditions used in this work lead to loss of BaP from solution (e.g., degradation, sorption to the container wall or bacteria), resulting in a non-detectable final amount by the time of measurement (Guo and Wen, 2021). For comparison, toxicokinetic data have been published for phenanthrene, structurally relatively similar to BaP, in *C. elegans* (Spann et al., 2015), using an exposure paradigm quite similar to ours except in chemically sterilized worms. As expected for a lipophilic chemical, phenanthrene bioaccumulated in the worms. Furthermore, the total concentration recovered was also a fraction of the original working concentration calculated at the start of exposure.

**Figure 2:**
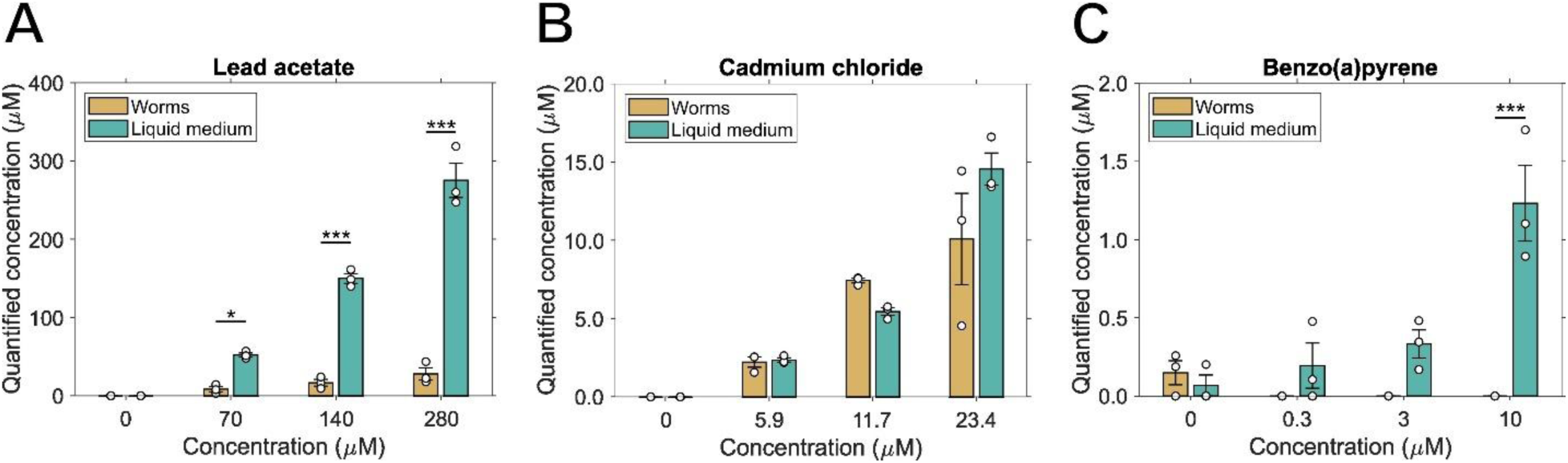
Chemical uptake of Pb, Cd, and BaP in *C. elegans*. Worms were exposed to multiple concentrations of Pb. Cd, and BaP, and after 24 hours these chemicals were quantified in both the worms and the supernatant from the liquid culture for (A) lead acetate, (B) cadmium chloride, and (C) benzo(a)pyrene. Two-way ANOVA with Tukey HSD, (*) *p* < 0.05, (***) *p* < 0.001.

### 3.3. Developmental exposures to Pb, Cd, and BaP cause morphological alterations in multiple neuronal types in late development (L4 stage)

Using the exposure protocol described above, we first asked whether we would encounter worms in which we would fail to identify the expected GFP-expressing neurons, when examined at the L4 stage that corresponds to complete development of their somatic cells including all neurons. For example, if a developmental exposure were to change the specification of a neuron from cholinergic to dopaminergic, we would expect to encounter cholinergic reporter worms in which some cholinergic neurons were absent, or dopaminergic reporter worms with extra, out-of-place DAT-1::GFP-expressing neurons. We never observed either.

Next, we tested whether the developmental exposures to Pb, Cd, and BaP would alter neuronal morphologies, assessed at the L4 stage. We used a previously developed scoring system (Bijwadia et al., 2021) to quantify alterations to the cephalic dopaminergic neurons (**Fig. 3A**). We developed similar scoring systems for glutamatergic, cholinergic, GABAergic, and serotonergic neurons as well as glial cells located in the head of *C. elegans* (**Fig. S2**). We observed an increase in morphological abnormalities after developmental exposure to Pb and Cd compared to the vehicle control for dopaminergic neurons (**Fig. 3A**). Glutamatergic neurons exhibited various degrees of morphological change after exposure to Pb, Cd, and BaP (**Fig. 3B**). In the case of cholinergic neurons, only exposure to BaP caused augmented changes (**Fig. 3C**). However, no significant differences from the vehicle control were observed for GABAergic neurons, serotonergic neurons, or for glial cells located in the head of *C. elegans* (**Fig. 3D-F**). Therefore, given the lack of response of these neurons to exposure to these classic developmental neurotoxicants, we performed further tests only with dopaminergic, glutamatergic, and cholinergic neurons. As a secondary benefit, the neurons of these types that we chose to score have very similar dendritic morphology and are all located close to each other in the head of *C. elegans*. For that reason, comparisons of the effects of a given exposure on morphology of these neuron types are more feasible both in terms of similarity to normal neuronal architecture, and in terms of toxicokinetic similarities.

**Figure 3:**
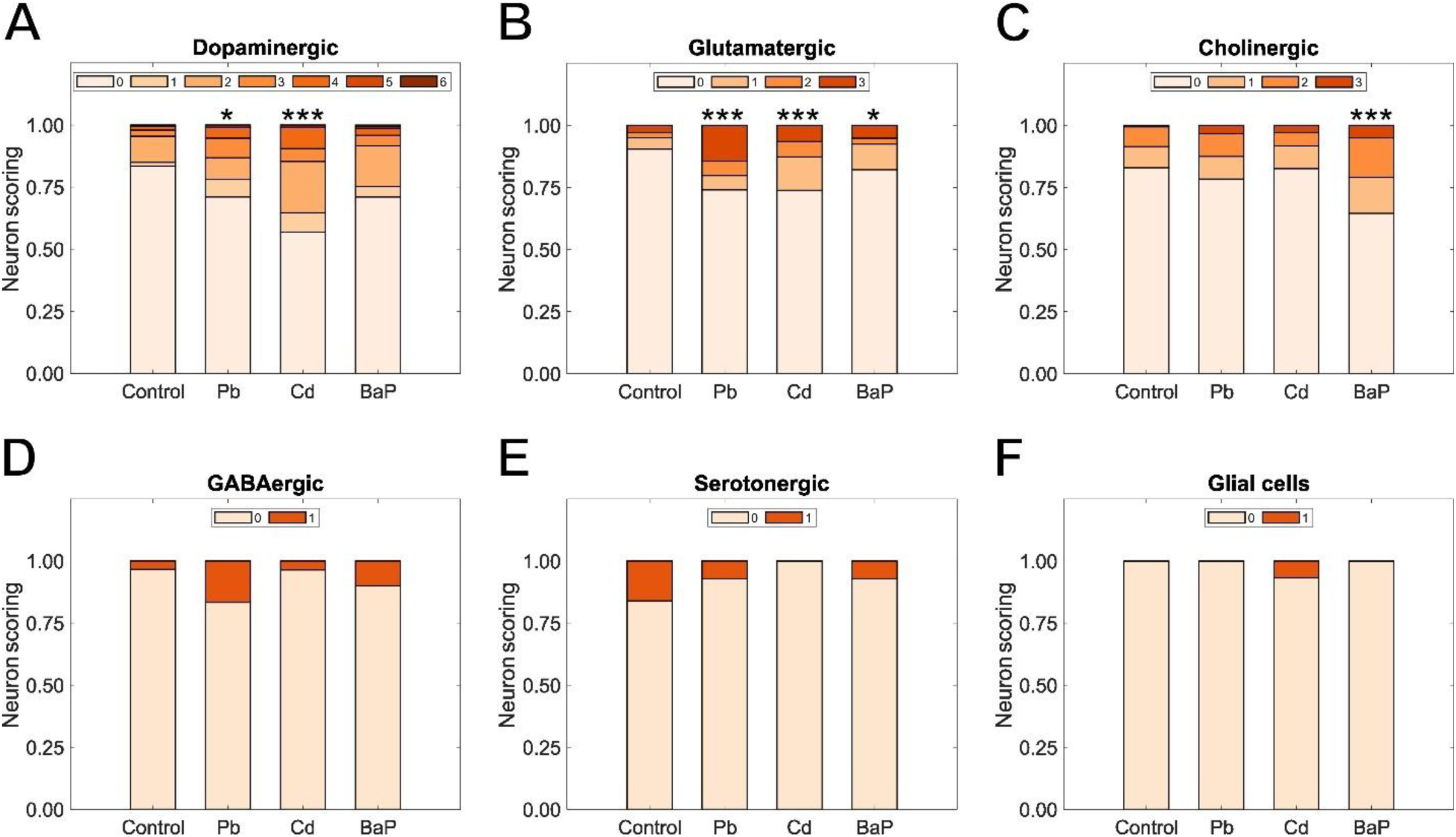
Neurodegenerative effects of developmental exposure to Pb, Cd, and BaP. Neuronal scoring in a 0-6 scale for (A) dopaminergic neurons; in a 0-3 scale for (B) glutamatergic and (C) cholinergic neurons; and in a 0-1 scale for (D) GABAergic neurons, (E) serotonergic neurons, and (F) glial cells. Chi-square test with Bonferroni correction, (*) *p* < 0.05, (***) *p* < 0.001.

### 3.4. Effects of secondary challenges to specific neurotoxicants in mid-adulthood

It is also possible that developmental exposure may cause neurotoxic effects that are only manifest later in life, or upon subsequent stress. Therefore, we assessed whether the developmentally exposed worms would show loss of neuronal integrity in day 4 of adulthood, with or without secondary challenge. For the dopaminergic neurons, we used the well-characterized dopaminergic neurotoxicant 6-hydroxydopamine (6-OHDA) (Nass et al., 2002). For cholinergic neurons, we used both chlorpyrifos, an organophosphate that inhibits acetylcholinesterase (Dam et al., 1999; Slotkin et al., 2001), and ethylcholine mustard aziridinium (AF64A), previously shown to produce cholinergic hypofunction in mice and rats (Fisher et al., 1982; Walsh et al., 1984). For glutamatergic neurons, we used quinolinic acid, reported to induce glutamatergic neurodegeneration in *C. elegans* (da Silveira et al., 2018). We determined experimental EC_50_ values based on morphological alterations as follows: 60.5 mM for 6-OHDA, 13.7 µM for chlorpyrifos, 77.0 µM for AF64A, and 94.1 mM for quinolinic acid (**Fig. S3**). This allowed us to test whether developmental exposures would alter subsequent EC_50_ values for morphological alterations.

We then took populations exposed during development to Pb, Cd, or BaP and subjected them to secondary challenge with the neurotoxicant matching its affected neuronal type at day 4 of adulthood. We observed a decrease in neurodegeneration in day 4 control worms (developmentally exposed to Pb, Cd, or BaP) in comparison to their L4 equivalents. This difference could be explained by neuron regeneration occurring during the 4 days between measurements (Wu et al., 2007). We found that worms exposed to 6-hydroxydopamine showed increased dopaminergic neurodegeneration when compared to their respective controls (**Fig. 4A**). This was the case for all developmentally exposed populations and for the control population. We observed that a challenge with quinolinic acid induced significant damage in glutamatergic neurons located in the head of worms for all developmentally exposed populations and controls (**Fig. 4B**). We also identified increased neurodegeneration in cholinergic neurons located in the head of worms challenged with chlorpyrifos with respect to non-challenged controls, but only for the control population and those worms exposed to Pb during development (**Fig. 4C**). Similarly for worms challenged with AF64A, we found that the population developmentally exposed to Pb, and the control population showed increased neurodegeneration in cholinergic neurons, but not those exposed to Cd or BaP (**Fig. 4D**).

**Figure 4:**
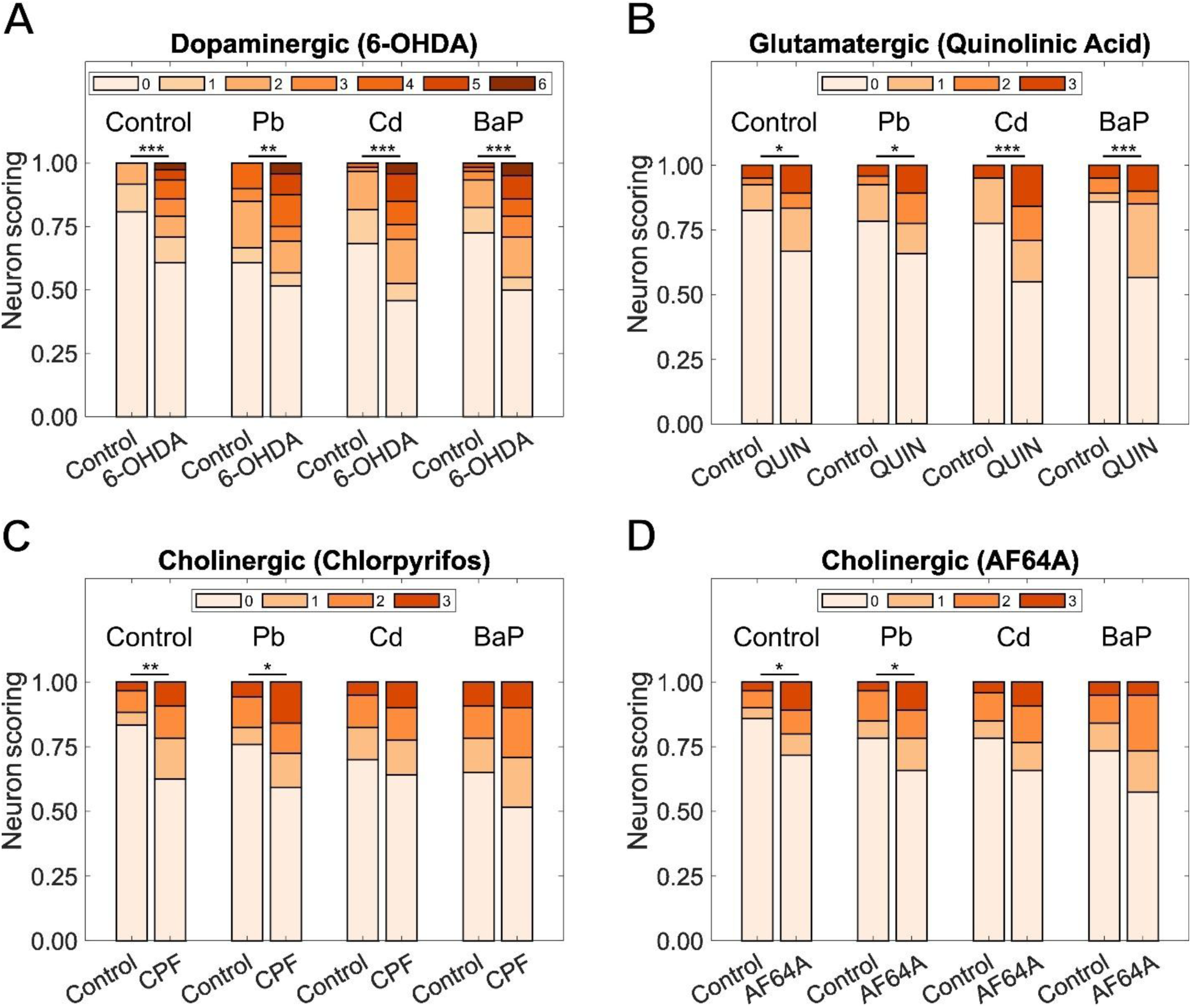
Effects of secondary challenges to selected neurotoxicants. Effects on different neuronal types after challenge to neuron-specific toxicants on day 4 of adulthood in worms developmentally exposed to Pb, Cd, and BaP: (A) 6-OHDA effect on dopaminergic neurons, (B) quinolinic acid effect on glutamatergic neurons, (C) chlorpyrifos and (D) AF64A effects on cholinergic neurons. Chi-square test with Bonferroni correction, (*) *p* <0.05, (**) *p* <0.01, (***) *p* <0.001.

### 3.5. Neuronal functional changes of developmentally exposed *C. elegans* in mid-adulthood

Next, to assess a functional rather than morphological alteration, we evaluated whether these developmental exposures might cause behavioral changes during mid-adulthood. We decided to measure behavioral responses related to each of the neuronal types we used for developmental exposures. Basal slowing response is a *C. elegans* behavior in response to the presence of bacterial food. Worms move slower in bacteria than in the absence of it to increase the time they spend in the presence of food (Sawin et al., 2000). This behavior, governed by dopamine signaling (Omura et al., 2012), is reported to be altered by exposure to Pb (Albrecht et al., 2022) and by morphological changes in dopaminergic neurons (Clark et al., 2024). Aldicarb is a carbamate that prevents the breakdown of acetylcholinesterase in the synapse cleft leading to paralysis in *C. elegans* (Opperman and Chang, 1991). Therefore, worms with defective or enhanced released of acetylcholine have different responses to Aldicarb compared to wildtype (Mahoney et al., 2006). Sensitivity to gentle touch in *C. elegans* is sensed by a set of six neurons (Goodman and Sengupta, 2019), and animals will perform forwards or backwards locomotion when touched in the tail or head respectively (Chalfie and Sulston, 1981). This behavior was altered in *C. elegans* after Aβ-induced neurodegeneration of glutamatergic neurons, and after exposure to the glutamatergic neurotoxicant quinolinic acid (Caldwell et al., 2020; da Silveira et al., 2018; Griffin et al., 2019). We allowed all developmentally exposed populations to reach day four of adulthood before proceeding with behavioral and neuronal function assays.

We assessed the effects of developmental exposures on the basal slowing response by measuring the number of body bends of *C. elegans* in plates containing bacterial food and without it (**Fig. 5A**). We tested positive control worms with mutations in the *cat-2* and *cat-4* genes. These dopamine deficient mutants are also defective for the basal slowing response (Omura et al., 2012; Sawin et al., 2000). Control worms show the expected reduction in movement on bacteria compared to empty plates. We observed no changes in response in worms exposed to BaP, but *C. elegans* exposed to Pb or Cd exhibited reduced basal slowing response.

**Figure 5:**
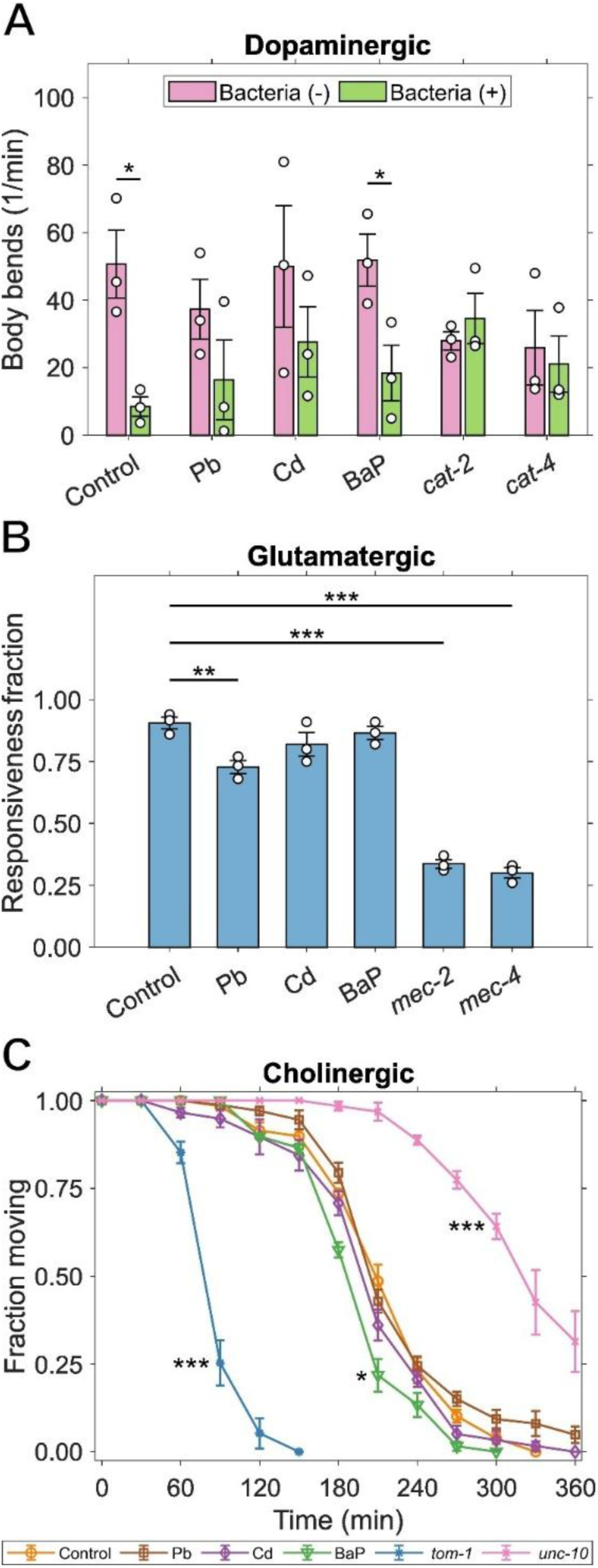
Effects of developmental exposure on neuronal functions during mid-adulthood. (A) Basal slowing response of worms developmentally exposed to Pb, Cd, or BaP; *cat-2* and *cat-4* mutants served as positive controls. (B) Time required to induce paralysis on Aldicarb plates for worms developmentally exposed to Pb, Cd, or BaP; *tom-1* and *unc-10* mutants served as positive controls. (C) Fraction of worms developmentally exposed to Pb, Cd, or BaP that are responsive to soft touch; *mec-2* and *mec-4* mutants served as positive controls. Two-way ANOVA with Tukey HSD for (A), One-way ANOVA with Dunnet’s test for (B), log-rank test and chi-square with Bonferroni correction for (C), (*) *p* < 0.05, (**) *p* <0.01, (***) *p* <0.001.

To measure the extent of behavioral changes caused by alterations in glutamatergic neurons, we tested the response of worms to soft touch by consecutively touching their heads and tails with an eyelash pick (**Fig. 5B**). We used positive controls with mutations in *mec-2* and *mec-4*, genes encoding elements required for mechanosensation channels (Goodman et al., 2002; Suzuki et al., 2003). We observed a moderate loss of sensitivity to soft touch in worms developmentally exposed to Pb, but not in those exposed to Cd or BaP.

We evaluated changes in cholinergic function by measuring the sensitivity to Aldicarb-induced paralysis in developmentally exposed worms (**Fig. 5C**). We used positive controls with mutations in the *tom-1* and *unc-10* genes, providing strong hypersensitivity and resistance to paralysis, respectively. These worms manifest altered synaptic transmission (Gracheva et al., 2006; Koushika et al., 2001) allowing for evaluation of the degree of Aldicarb resistance or sensitivity in treatment worms. We did not find any differences in the onset of paralysis for worms treated with Pb or Cd compared to control. However, we observed that those treated with BaP had a slight increase in sensitivity to Aldicarb.

## Discussion

We present a DNT assay that will allow researchers to leverage the many strengths of *C. elegans* in DNT research. An important improvement compared to previous DNT assays is an exposure paradigm that includes most or all neurodevelopmental processes. Major improvements compared to previously described general neurotoxicological/neurodegenerative assays include examination of multiple neuron types, including chemical and genetic positive controls for morphological and behavioral assays for dopaminergic, cholinergic, and glutamatergic neurons. We found that exposure to Pb, Cd, and BaP cause alterations to neuronal structure and function that are generally comparable to those previously reported in vertebrate models, with some novel observations that merit further experimentation. Finally, we did not find exposures that resulted in conversion of one type (defined by strains designed to report neurotransmitter type) of neuron to another. We discuss this and other limitations and uncertainties associated with the *C. elegans* DNT model.

We designed a DNT exposure protocol that incorporates developmental exposures on gravid worms of the parent generation to simulate *in utero* exposure that is known to be important for these pollutants (Chandravanshi et al., 2021; Lin et al., 2013; Meier et al., 2017; Ortiz et al., 2013). In this manner, we aimed at measuring effects caused by exposures throughout embryonic development, encompassing most or all of neurodevelopment. As described, we think it is important to use concentrations high enough to cause enough biological effect that lack of DNT is unlikely to be a false negative, but low enough that DNT outcomes are unlikely to be secondary to overt organismal toxicity. However, the hypodermis of *C. elegans* has been proven to act as a barrier for the uptake of some chemicals present in their culturing media (Xiong et al., 2017), making it very challenging to relate media concentrations of toxicants used in C. elegans experiments to those observed in people (e.g., blood concentrations) or used in cell culture experiments. Therefore, we performed quantification of internal uptake of Pb, Cd, and BaP. We found for Pb that internal concentration in worms was 11% to 16% that of the media, and for cadmium chloride it was at least 73% that of the media. A fraction of these chemicals was present inside of the worms’ bodies, indicating that the neurotoxic effects we observed occurred at lower concentrations than those measured in the worms’ media. We found that the internal concentrations in exposed worms were 343 µg/dL for Pb, 84 µg/dL for Cd, and below the detection limit (0.98 µg/dL) for BaP. Previously reported values for human blood concentrations range from 5 – 30 µg/dL for Pb, 0.5 – 1 µg/dL for Cd, and 50 – 500 fg/dL for BaP with the caveat that BaP is rapidly eliminated with a half-life averaging 46.5 hours (Edwards et al., 2015; King et al., 2015; McKelvey et al., 2007; Vermillion Maier et al., 2022), indicating that internal concentration in worms were higher than those found in human blood by approximately one order of magnitude. However, studies in human and animal cell models with these chemicals used ranges of 0.08 – 50 µM for Pb, 40 – 60 µM for Cd, and 0.25 – 5.0 µM for BaP, which are close to our internal concentrations of 16.6 µM for Pb, 7.4 µM for Cd, and below limit of detection (0.039 µM) for BaP (Gillis et al., 2012; Hockley et al., 2006; Olabarrieta et al., 2001).

Different neurotoxicants target different cell types, for reasons that are not often well understood. We measured neurodegeneration for all neuronal types at the last larval stage of *C. elegans* because at this stage all of its somatic cells would be fully developed, permitting using the same scoring systems and comparison with older worms. We observed that the neurotoxic effects of Pb, Cd, and BaP were distinct and dependent on neuronal type, with only dopaminergic, cholinergic, and glutamatergic neurons showing any significant amount of damage. As an added benefit from the perspective of directly comparing the effects of exposure on different neuron types, these three neuronal types share similar structural morphology in the head area of *C. elegans*. Their cell body is located just before the end of the pharynx and start of the intestine, with dendrites extending from there to the tip of the worm’s head. Thus, these neuron types can be compared for sensitivity to different toxicants while excluding the possible influence of large morphological differences (e.g., shape and size) and intraorganismal toxicokinetic differences (although cell-specific toxicokinetic differences such as specific transporter expression remain).

Dopaminergic, cholinergic, and glutamatergic neurons differed in their sensitivity to chemical-induced damage at the low-toxicity concentrations used in this work. Dopaminergic neurons were affected by heavy metals Pb and Cd, but not by the polycyclic aromatic hydrocarbon (PAH) BaP. Inversely, only BaP caused significant damage to cholinergic neurons, while glutamatergic neurons suffered effects from all exposures. There is evidence that increase in production of reactive oxygen species, expected to be induced by Pb or Cd, is the leading cause of the observed DNT (Leal et al., 2012; Struzyńska, 2009; Vellingiri et al., 2022). Cholinergic neurons may be less sensitive to these effects but more likely to be affected by signaling disruption from exposure to BaP (Chepelev et al., 2015; Slotkin et al., 2019). However, it is important to note that the worm homolog to the aryl hydrocarbon receptor, a major mediator of BaP signaling in vertebrates, may not be activated by BaP (Karengera et al., 2022; Powell-Coffman et al., 1998), and worms lack the ability to metabolize BaP to DNA-reactive forms (Leung et al., 2010). The likelihood that BaP does not interact with worm AHR-1 is supported by the lack of effect of BaP on GABAergic neurons, despite the known role for AHR-1 in GABAergic neuron development in *C. elegans* (Huang et al., 2004).

Our observation of lead and cadmium causing dopaminergic neurodegeneration in *C. elegans* is consistent with previous work showing that worms exposed to lead exhibit altered dopaminergic morphology and impairment in neuronal function tested by BSR (Akinyemi et al., 2019; Albrecht et al., 2022), and that both lead and cadmium exposure induced abnormalities in dopaminergic neurons, paired with decreased body length, brood size and feeding (Tang et al., 2019). Similarly, rats maternally and permanently exposed to lead exhibited higher locomotor activity connected to the cortical dopaminergic system (Ma et al., 1999), prenatal exposure to lead in pregnant rats accentuated behavioral effects in the offspring related to increased dopamine turnover in basal ganglia (Szczerbak et al., 2007), and prenatal exposure to lead in rats caused functional dopamine system changes (Stansfield et al., 2015). In rats, cadmium caused excessive dopamine release from striatal slices (Gutiérrez-Reyes et al., 1998), decreased ^3^H-dopamine uptake indicating sensitivity of dopaminergic neurons to cadmium toxicity (Yang et al., 2007), and decreased dopamine in the median eminence while increasing it in the posterior pituitary (Lafuente et al., 2005). Reported effects of Pb, Cd, and BaP in glutamatergic neurons in *C. elegans* are scarce, but exposure to diesel exhaust particles containing PAHs lead to degeneration in glutamatergic neurons in *C. elegans* (Chatterjee et al., 2024). However, it has been reported that rats chronically exposed to lead have deficit in glutamatergic neurotransmission (Neal and Guilarte, 2013), and gestational exposure to BaP in rats caused down-regulation of glutamatergic receptor unit expression (Brown et al., 2007). Although there are no reports of the effects of BaP in cholinergic neurons using *C. elegans*, developmental exposure to BaP on rats impaired acetylcholine presynaptic activity (Slotkin et al., 2019).

Later-life consequences, such as neurodegeneration or increased vulnerability to stressors, are important yet still poorly understood manifestations of the Developmental Origins of Health and Disease (Gao et al., 2024; Miller and O’Callaghan, 2008), or DOHAD. Therefore, we moved on to evaluating the later-life effects of these developmental exposures. We looked at resistance to neurotoxicant challenges (assessed morphologically), and functional changes in *C. elegans* during mid-adulthood (without secondary challenge). We did not observe any increase or decrease in resilience to secondary challenge with neuron-specific neurotoxicants except for Cd and BaP in cholinergic neurons. In these two cases, we measured no statistically significant differences with their respective control populations, indicating that the pre-exposure was protective against subsequent challenge. However, both manifest the same trend towards increased neurodegeneration in challenged populations as in all the other cases. It is possible that instead of a change in resilience, what we observed is the effect of increased baseline neurodegeneration in the control populations caused by previous exposure or increased age. In general, early life exposures can cause either increased resiliency (sometimes referred to as hormesis) or decreased resiliency (DOHAD) later in life.

However, we did observe many behavioral changes in mid-adulthood, and these corresponded very well to the neuronal types that showed significant morphological damage after development. We evaluated these changes during mid-adulthood and found that even after 4 days of adulthood, worms were still affected by their early exposures. The basal slowing response, regulated by dopaminergic signaling, was lost in worms exposed to Pb and Cd. Notably, these two chemicals also caused neurodegeneration measured at the L4 larval stage. Similarly, BaP was the only chemical that induced damage in cholinergic neurons when measured at the L4 larval stage, and the only exposure leading to increased sensitivity to Aldicarb-induced paralysis. These results support a direct relationship between DNT and later-life behavioral changes. However, for glutamatergic neurons, we observed loss of mechanosensory response only for worms exposed to Pb, but we originally observed neuronal damage for all three chemicals evaluated. A possible explanation for this discrepancy is that we needed to reach a threshold for early neurodegeneration to obtain changes in behavior, as suggested by the fact that Cd induced a trend towards reduced mechanosensation, but this was not statistically significant.

Despite these encouraging results, the simplicity of the *C. elegans* model does impose some limitations. For example, it may limit our ability to detect effects that require complex physiological interactions (e.g., microbiome-vagus nerve-dopaminergic neurons) (Braak et al., 2003; Klingelhoefer and Reichmann, 2015), although nerve-nerve transmission of transgenically expressed α–synuclein has been observed (Tyson et al., 2017) and gut-neuron interactions (Chikka et al., 2016) as well as glial cell-neuron interactions (Gibson et al., 2018) have both been studied in *C. elegans*. In addition, *C. elegans* lacks genes encoding for voltage-gated sodium channels (Gao and Zhen, 2011), leading to its use of calcium channels to generate action potentials (Liu et al., 2018), suggesting that neuronal electric signaling in *C. elegans* spreads passively rather than having an active propagation (Goodman et al., 1998). Also, *C. elegans* does not exhibit axonal myelination (Oikonomou and Shaham, 2011), with damage to the myelin sheath identified as a contributor to impaired neuronal function (Llorens et al., 2011). We focused on looking at neurons under one neurotransmitter at a time, but some neurons in *C. elegans* express multiple neurotransmitters (Etchberger et al., 2007; Poole et al., 2024), making our approach imperfect. Finally, our work suggests that while genetic mutations can lead to loss or gain of specific neuronal subtypes, the process of neuronal specification may be so hardwired that it is quite robust to chemical exposure, limiting the utility of this model to detect DNT that takes the form of changed neuronal specification.

Future directions should include using our approach to expand to other human DNT chemicals, particularly those that could affect the neuronal types that were not damaged by exposure to lead, cadmium or BaP. For instance, nickel has been shown to induce GABAergic neurodegeneration (Ijomone et al., 2020), and manganese altered GABA metabolism in rat brains (Erikson and Aschner, 2003); parathion and diazinon caused altered serotonin function in rats (Slotkin et al., 2009, 2008); and an As-Cd-Pb mixture induced reduction in glial fibrillary acidic protein during brain development of rats (Rai et al., 2010). This suggests that metals other than Pb and Cd could cause DNT in GABAergic neurons, organophosphates could induce DNT in serotonergic neurons, and glial cells may require a mixture design to elicit DNT. Our model could also be used to test DNT of as-yet untested chemicals and mixtures. It could be improved still further by examining potential effects of chemical exposure on synapse formation, another process that is nearly invariant and thus relatively easy to study for deviations from normal development. Furthermore, using a worm with all neurons identified following a fluorescent color map (Yemini et al., 2021) could facilitate faster association of chemical-induced DNT with specific neuronal types. Importantly, the use of our DNT experimental approach using *C. elegans* could serve to increase the confidence in new approach methodologies (NAM) by serving as a bridge in the middle between *in vitro* systems and vertebrate models (van der Zalm et al., 2022). This is highlighted by the ability to observe cell-cell interactions (Singhvi and Shaham, 2019); the effects of other factors such as exercise (Laranjeiro et al., 2019), diet (Bishop and Guarente, 2007), and temperature (Prahlad et al., 2008; Tatum et al., 2015); and the ability to perform cost-effective drug-screening focused on neurodevelopmental and neurodegenerative therapeutics (Romussi et al., 2024). Finally, we consider that it is important in these studies to continue to quantify biological uptake of chemicals and consider speciation and partitioning of chemicals in exposure chambers. This is critical both to understanding internal concentrations for the purpose of comparing to other studies, and to understand the limitations that may occur with chemicals with low water solubility, high volatility, or other properties that limit bioavailability.

## Conclusion

In this work, we developed an experimental approach for DNT assessment of the full developmental timecourse, examining both morphological and functional neurotoxicological effects during mid-adulthood in the nematode *C. elegans*. Our approach was developed with the well-studied developmental neurotoxicants Pb, Cd, and BaP, but can be applied to other toxic chemicals. An advantage of this model is the unambiguous separation of structural versus functional forms of DNT in different neuronal types. DNT effects can be driven both by long-term alterations in cell fate and architecture (“hardwiring”) (Di Consiglio et al., 2020; Slotkin et al., 2016), as well as long-term and persistent functional changes in cells that are apparently not altered in cell type (“programming”) (Wan et al., 2021; Zhou et al., 2020), although these can be difficult to distinguish and are not mutually exclusive. Indeed, it is likely that many exposures are likely to alter both hardwiring and programming. Our experimental platform enables determination of late-life effects of DNT for multiple neuronal types in an intact and exceptionally developmentally and neurologically well-studied organism, reducing the need for more expensive vertebrate models and creating a bridge between those models and other NAMs, ultimately increasing the speed of DNT data acquisition to evaluate the risk of chemical exposure.

## Credit authorship contribution statement

Javier Huayta: investigation, methodology, formal analysis, writing-original draft, visualization. Sarah Seay: investigation, methodology, formal analysis. Joseph Laster: investigation, formal analysis. Nelson A. Rivera Jr.: investigation, methodology, formal analysis. Abigail S. Joyce: investigation, methodology, formal analysis. Patrick L. Ferguson: resources. Heileen Hsu-Kim: resources. Joel N. Meyer: conceptualization, funding acquisition, methodology, project administration, writing original-draft, supervision.

## Acknowledgements

Some strains were provided by the CGC, which is funded by NIH Office of Research Infrastructure Programs (P40 OD010440). c-elegans-egg_laying-defective icon, c-elegans_small_mutant icon, 24-well_plate icon, and on_Nematode_Growth_Medium_petri_plate icon by DBCLS https://togotv.dbcls.jp/en/pics.html is licensed under CC-BY 4.0 Unported https://creativecommons.org/licenses/by/4.0/

## Data availability

All data in this study is included in this article, its supplementary information flies, and it is available from the corresponding author upon request.

