## Supplemental figures for "Assessment of developmental neurotoxicology-associated alterations in neuronal architecture and function using *Caenorhabditis elegans*"

**Supplementary information**

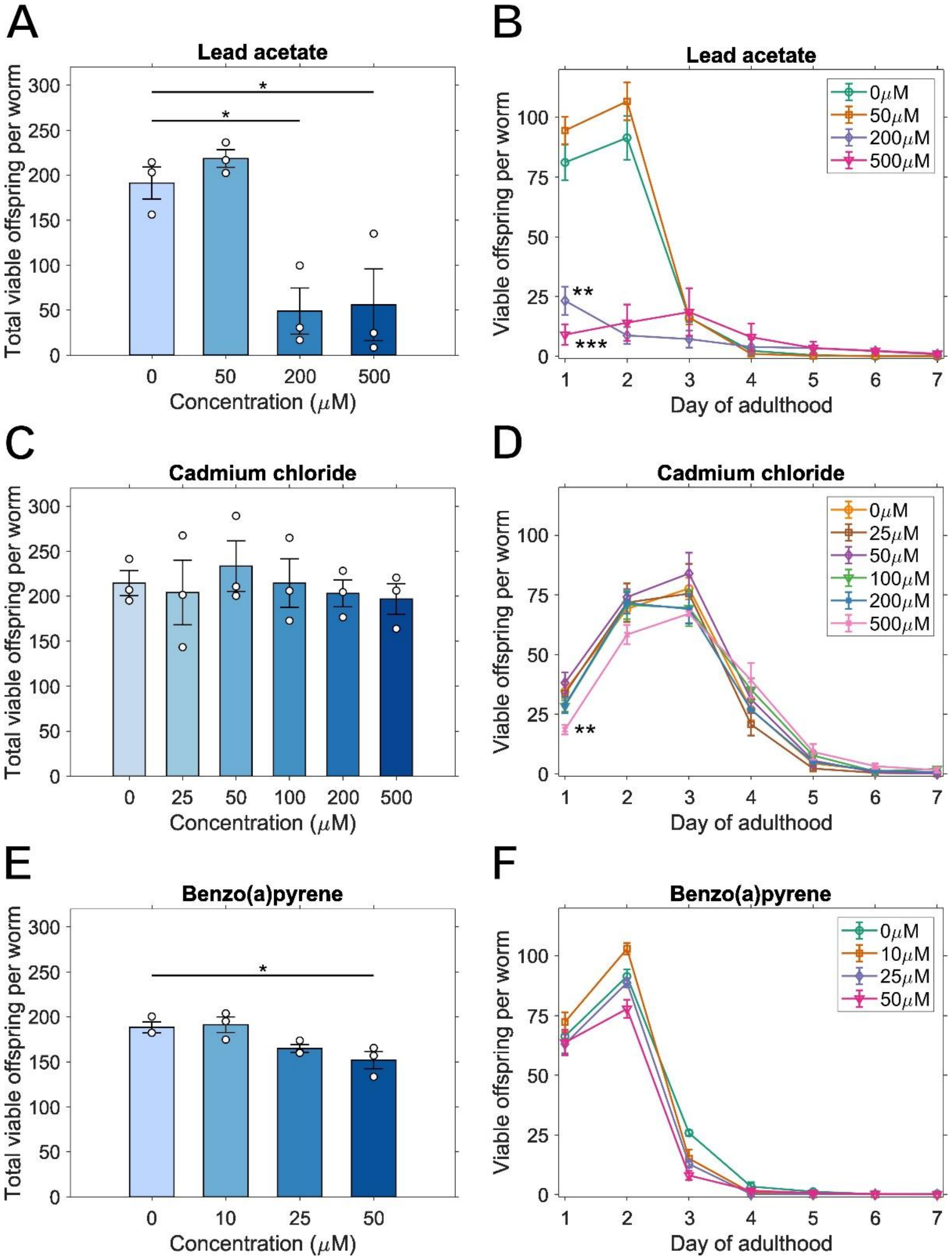


**Figure S1:** Quantification of effect on reproduction. Effects of lead exposure on (A) total viable offspring and (B) viable offspring per day. Effects of cadmium exposure on (C) total viable offspring and (D) viable offspring per day. Effects of benzo(a)pyrene exposure on (E) total viable offspring and (F) viable offspring per day. One-way ANOVA with Dunnet's test for (A), (C), and (E), log-rank test and chi-square with Bonferroni correction for (B), (D), and (F), (*) $p < 0.05$, (**) $p < 0.01$, (***) $p < 0.001$.

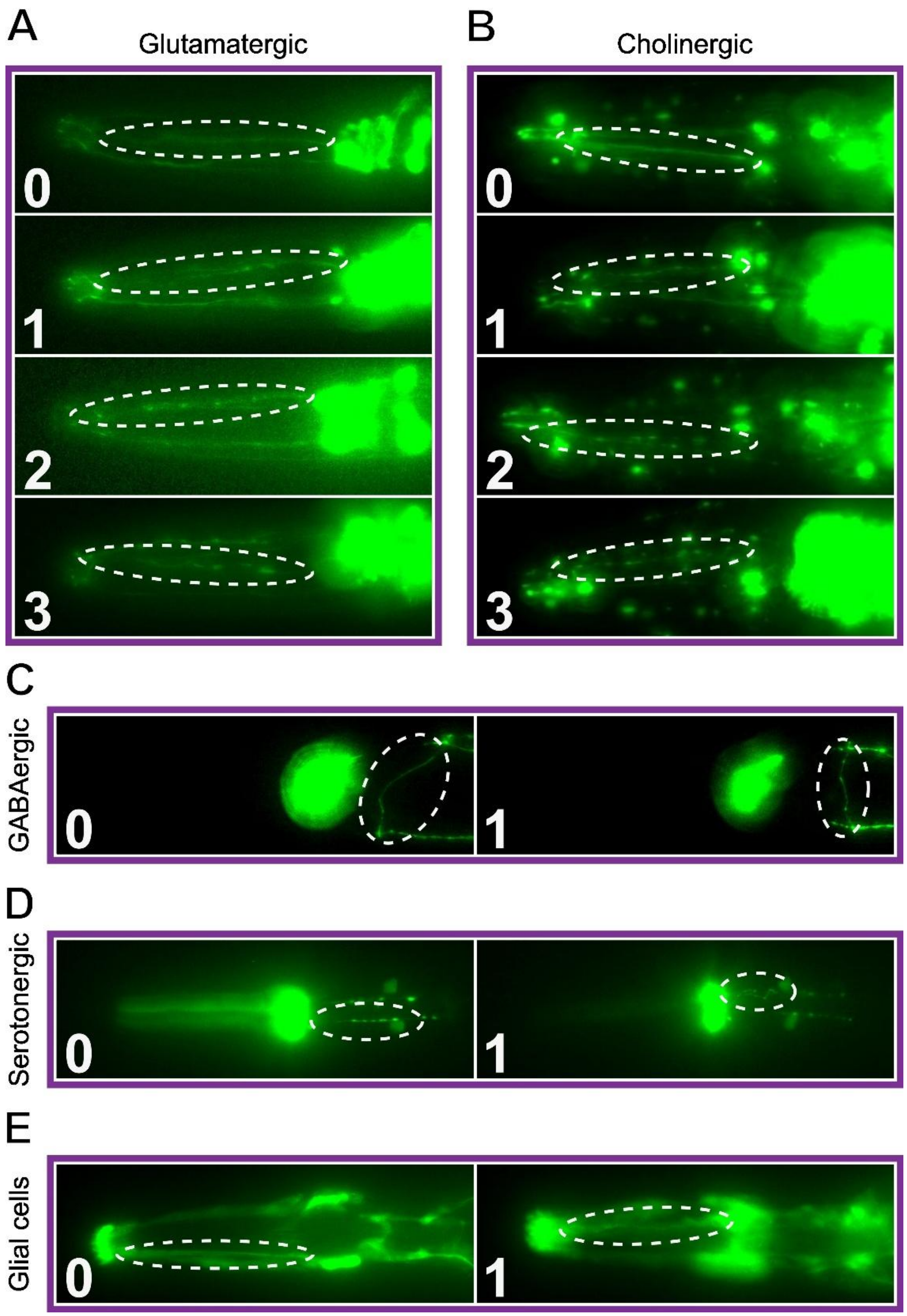


**Figure S2:** Scoring systems for quantification of neuron damage. (A) Glutamatergic neurons: 0 – no damage, 1 – 1 to 5 blebs, 2 – 6 to 10 blebs, and 3 – more than 10 blebs and/or presence of breaks. (B) Cholinergic neurons: 0 – no damage, 1 – irregular (curves), 2 – 1 to 10 blebs, and 3 – more than 10 blebs and/or presence of breaks. (C) GABAergic neurons: 0 – no damage, and 1 – any type of damage. (D) Serotonergic neurons: 0 – no damage, and 1 – any type of damage. (E) Glial cells: 0 – no damage, and 1 – any type of damage.

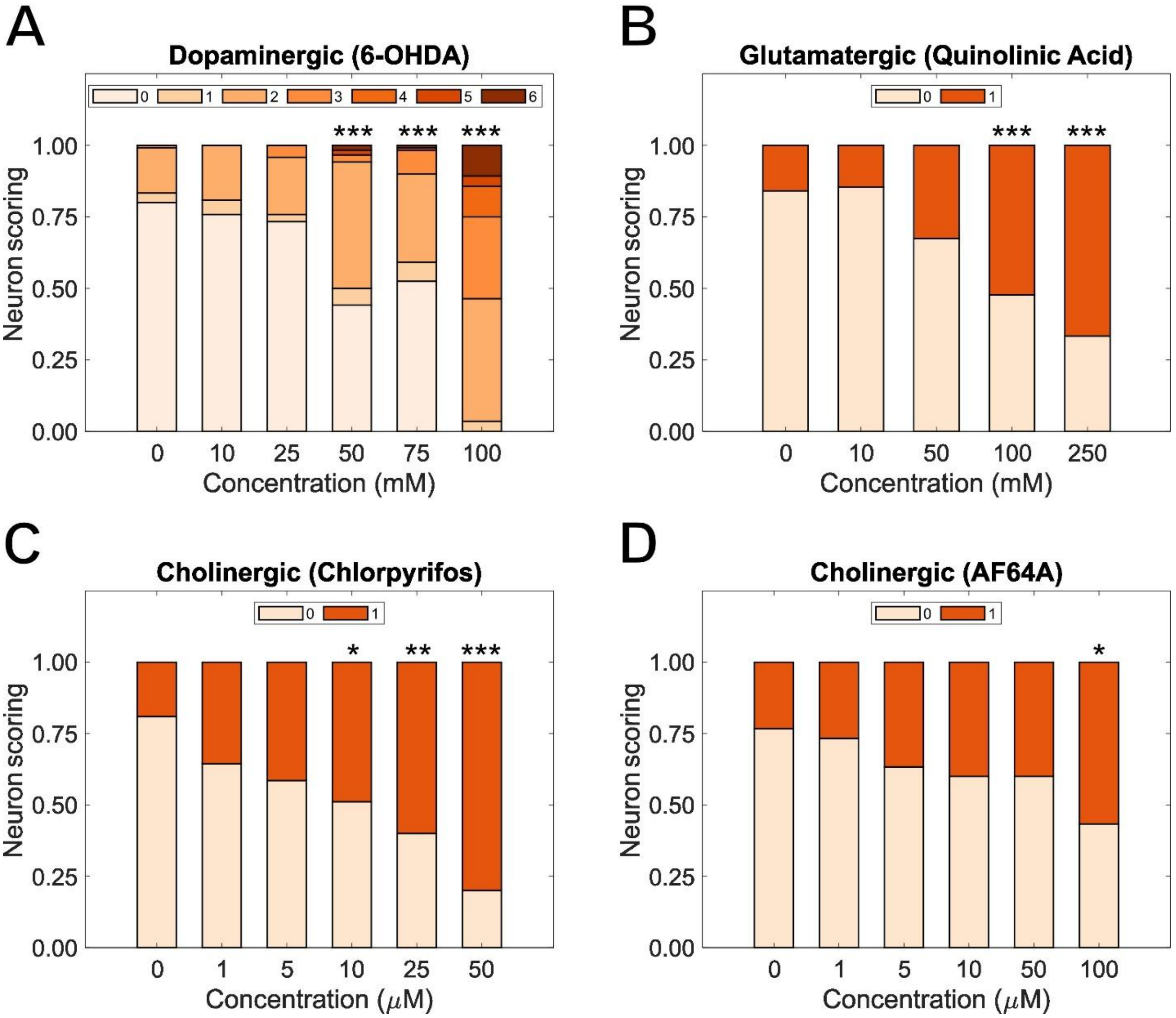


**Figure S3:** Determination of $EC_{50}$ values for neurodegeneration after exposure to neuron-specific neurotoxicants. (A) 6-hydroxydopamine for dopaminergic neurons. (B) Quinolinic acid for glutamatergic neurons. (C) Chlorpyrifos and (D) AF64A for cholinergic neurons. Chi-square test with Bonferroni correction, (*) $p < 0.05$, (**) $p < 0.01$, (***) $p < 0.001$.
